# IMMUNOLOGICAL INVESTIGATIONS OF HHV-6 U12 AND U51 ENCODED PROTEINS INVOLVEMENT IN AUTOIMMUNE THYROIDITIS DEVELOPMENT

**DOI:** 10.1101/2020.07.20.211698

**Authors:** Alina Sultanova, Maksims Cistjakovs, Liba Sokolovska, Egils Cunskis, Modra Murovska

## Abstract

Human herpesvirus 6 (HHV-6) is a human pathogen with a wide cell tropism and many immunomodulating properties. HHV-6 has been linked to the development of multiple diseases, among them – autoimmune. Conflicting evidence implicates HHV-6 in autoimmune thyroiditis (AIT). HHV-6 contains two genes (U12 and U51) that encode putative homologues of human G-protein-coupled receptors (GPCR) like CCR1, CCR3 and CCR5. It has been shown that proteins encoded by HHV-6 U12 and U51 genes can be expressed on the surface of epithelial and some peripheral blood mononuclear cells populations, which makes them a potential cause for evoking autoimmunity.

The aim of this study was to identify potentially immunogenic synthetic peptides derived from HHV-6 U12 and U51 amino acid sequences and to find evidences of the possible involvement of these proteins in AIT development. 62 AIT patients positive for HHV-6 infection were enrolled in this study. 30 different synthetic peptides designed from HHV-6 U12 and U51 proteins’ amino acid sequences, as well as, recombinant human CCR1, CCR3 and CCR5 proteins were used for suspension multiplex immunological assay (SMIA) to detect specific IgG, and IgM antibodies.

HHV-6 peptide specific IgG and IgM antibodies were found in patient’s samples, with higher signals for IgM antibodies, which is indicative of reactivation and active HHV-6 infection. As well recombinant CCR1 and CCR5 showed high signals on IgM antibodies which is indicating on the presence of potential auto-antibodies against human G protein-coupled receptors. No cross reactivity between HHV-6 peptide specific antibodies and human recombinant CCR1, CCR3 and CCR5 was found, however, the possibility of cross-reactive autoantibodies specific for structural epitopes cannot be excluded.

## Background

HHV-6 is a widely distributed human pathogen which can achieve lifelong persistence in its host after primary infection occurring in early childhood. Even though the pathogen was discovered more than 30 years ago, many aspects of its infection and its role in disease development are still poorly understood.

HHV-6 infection can affect the host in the long term and contribute to several autoimmune disorders, including autoimmune hemolytic anemia/neutropenia [1], autoimmune acute hepatitis [2], and multiple sclerosis [3–5]. Recently, more and more attention has been given to the possible involvement of viruses into the development of autoimmune thyroiditis.

The study published by Caselli et al. (2012) links HHV-6 to Hashimoto thyroiditis (HT). This study demonstrated that thyroid fine needle aspirates (FNA) obtained from patients with HT revealed the presence of HHV-6 significantly more frequently in comparison with the controls (82% and 10%, respectively). Furthermore, active HHV-6 transcription was observed in HT thyrocytes, compared with the latent infection found in the HHV-6 infected thyrocytes used as a control. These researchers proposed a potential mechanism for HHV-6-induced autoimmunity demonstrating that follicle cells infected with HHV-6 became susceptible to NK-mediated killing [6]. Also, our previously published data showed an almost 100% incidence rate of HHV-6 genomic sequence in thyroid gland tissue samples acquired from AIT patients’ post-surgical materials [7]. Furthermore, we found a significantly higher presence of HHV-6 activation marker (HHV-6 U79/80 mRNA) in AIT patients’ samples in comparison to control groups’ [18/44 (41%) vs. 1/17 (6%), respectively; p 0.0118] [7].

HHV-6 possesses a number of immunomodulating properties for immune evasion and viral dissemination. These include the ability to alter the repertoire of molecules expressed on cell surfaces and chemokine and cytokine expression and secretion [8]. Another immunomodulation strategy utilized by herpesviruses, including HHV-6, is the ability to encode viral chemokines and chemokine receptors [9].

HHV-6 encodes two viral chemokine receptors U12 and U51, which are structurally similar to cellular G protein-coupled receptors (GPCR) [10], but the role of these genes is not well understood. HHV-6 U12 and U51 are functionally similar to the much better studied CMV protein US28 (pUS28), which enhances the course of CMV infection. As a CCR homologue, pUS28 dimerizes with a number of host chemokine receptors, including CCR1, CCR2, CCR3, CCR4, CCR5, and CXCR4 [11]. HHV-6 GPCRs are not sufficiently studied, which raises the question of whether these proteins are involved in the development of the autoimmune processes and how it would affect the host’s immune system if it were able to dimerize with host chemokine receptors similar to the hCMV protein US28.

It has been shown that proteins which are encoded by HHV-6 U12 and U51 genes can be expressed on the surface of epithelial and some peripheral blood mononuclear cells populations, which makes them a potential cause for evoking autoimmunity by making hosts GPCRs targets for auto-reactive T and B lymphocytes [12,13]. Additionally, it has been shown *in vitro* that HHV-6 encoded GPCRs could interact with cytokine signalling pathways by down-regulating RANTES [14].

Knowing the structural and functional similarity of HHV-6 U12 and U51 proteins with cellular GPCRs, it is worth considering their role in autoimmunity development, but both HHV-6 encoded chemokine receptors are poorly studied and studies that have been conducted were done more than 20 years ago, which means their role in HHV-6 infection and disease development is unclear.

One of the reasons why HHV-6 U12 and U51 gene encoded proteins are poorly studied is because of the transmembrane nature of these proteins and the difficulty in purifying them in recombinant protein acquirement. To acquire antigens, which could be used effectively for immunisation and investigation of the possible involvement of these proteins in autoimmune diseases development, synthetic peptides from HHV-6 U12 and U51 amino acids sequences were designed in this study. Although, presentation of potential linear epitopes in form of synthetic peptides could not give the whole immunogenic response picture, it provides a much cheaper and faster approach in research of HHV-6 proteins.

The aim of this study was to identify potentially immunogenic synthetic peptides derived from HHV-6 U12 and U51 amino acid sequences and to find evidences of the possible involvement of these proteins in AIT development.

## Materials and methods

### Peptides and study group

Alignment of GPCR viral putative homologues (HHV-6A/B U12 and U51) and human (CCR1, CCR3 and CCR5) by T-coffee software, as well as, Bepipred Linear Epitope Prediction and Kolaskar & Tongaonkar algorithms were used for linear epitope prediction to design 20 mer synthetic peptides (Figure 1). All synthetic peptides were modified with three polyethylene glycol (PEG3) molecules at amino terminus end. PEG3 is used as a spacer in carbodiimide coupling of peptides to carboxylated magnetic beads, to provide better freedom to present epitopes to antibodies during SMIA protocol.

**Figure 1.**
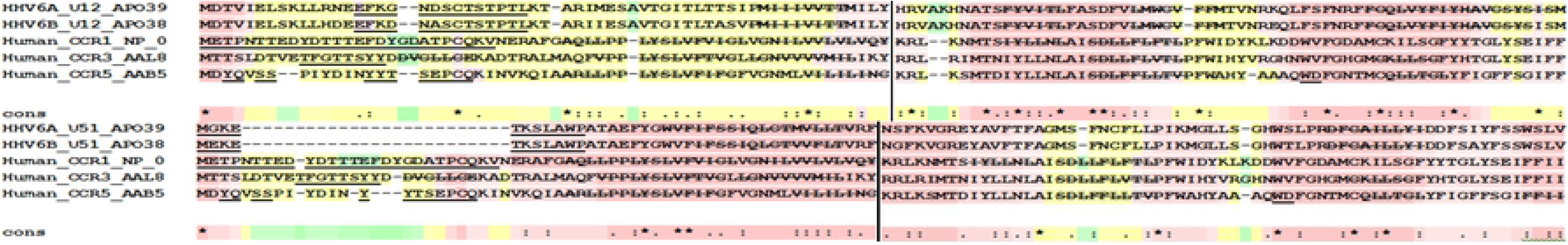
Example of high conservative (pink) regions’ alignment (region 1-126) and distribution of predicted linear epitopes (Bepipred Linear Epitope Prediction[underline] and Kolaskar & Tongaonkar [strikethrough]) for HHV-6A/B U12, HHV-6A/B U51, as well as, human CCR1, CCR3 and CCR5 genes.

HHV-6 U12 and U51 amino-acids sequences from both HHV-6 variants - A and B, were taken for peptide design. In total 30 synthetic peptides were designed (Table 1).

**Table 1.**
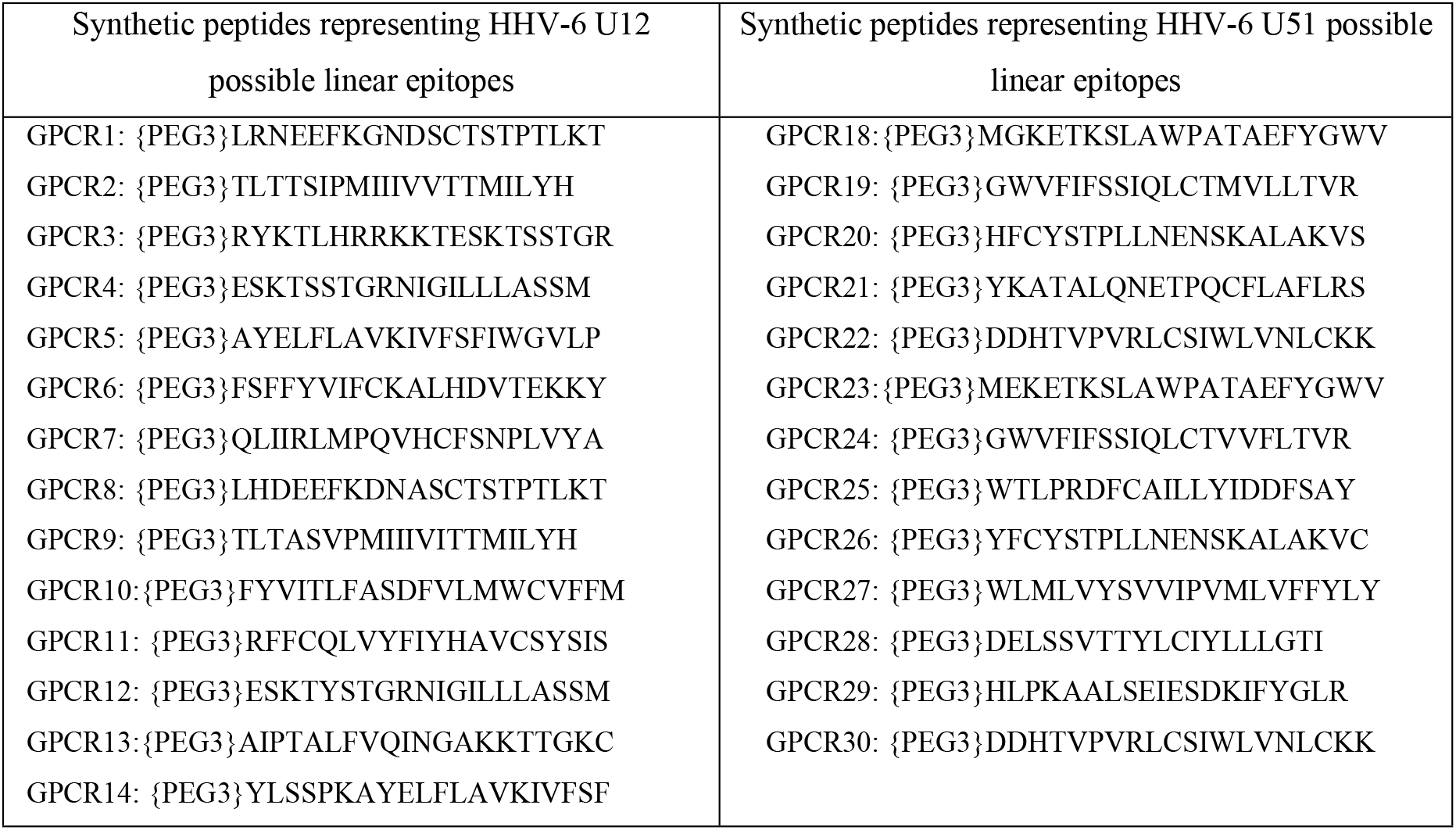
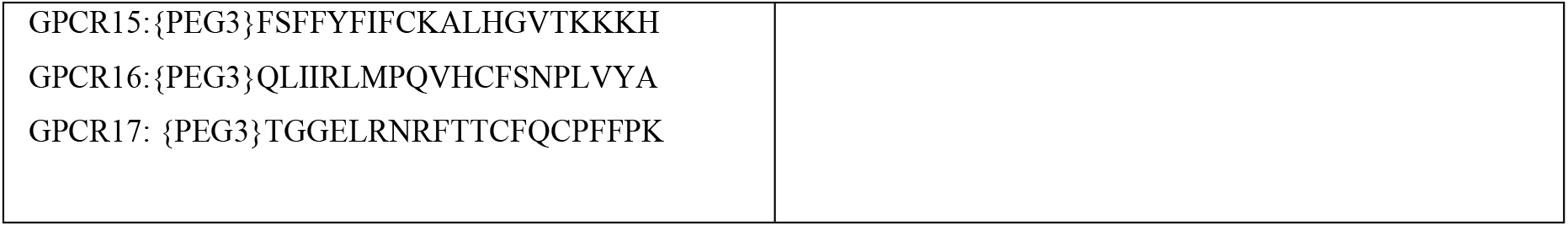
List of acquired synthetic peptides

The peptides were synthesized by GenScript. Lyophilized peptides were dissolved in sterile phosphate-buffered saline (PBS), pH 7. The dissolution sometimes had to be facilitated by warming the solution to 37°C overnight on a shaker or by sonication for 5 min. Highly hydrophobic peptides were dissolved in 10 to 50% (vol/vol) dimethyl sulfoxide (DMSO) (Sigma D2650).

Human recombinant CCR1, 3 and 5 were obtained from Abnova (Taipei, Taiwan). The proteins were produced in vitro using wheat germ expression system which preserves native protein folding.

62 patients with autoimmune thyroiditis following thyroidectomy positive for HHV-6 infection were enrolled in this study, of which 4 were males (6%) and 58 were females (94%), with median age of 52 (interquartile range [IQR]: 42-61).

The permission to conduct the research was received from the Riga Stradins University (RSU) Ethics Committee and all participants in the study gave their written consent to the examinations. Samples were received from the Riga East Clinical University Hospital.

### Coupling of proteins and peptides to magnetic carboxylated microspheres

The synthetic peptides or protein were coupled to Luminex Magnetic Carboxylated Microspheres (MagPlex) using xMAP^®^ Antibody Coupling Kit. The final pellet was resuspended in an appropriate amount of StabilGuard buffer (1000 μl of StabilGuard buffer if 100μl of beads were taken). This created a bead mixture consisting of 1250 beads/μl. The coupled beads were stored at 4°C in the dark.

### Suspension Multiplex Immunoassay (SMIA)

The assay system required only 5-10 μl of plasma. Antigen was coupled covalently to carboxylated color-coded beads as described above. IgG was detected using biotinylated protein G (BPG) as secondary antibody. For IgM antibody detection biotinylated anti-human IgM (affinity purified, μ-chain specific, Sigma-Aldrich cat. Nr. B1140) was used as secondary antibody.

The principal of SMIA is similar to ELISA - immunosorbent ‘sandwich’ type assay. However, reaction in SMIA runs in suspension, which gives better freedom for immunosorbent reaction, and sensitivity of such assay would always be more sensitive than any ELISA. In combination with Luminex 200 and MagPlex system, SMIA allows the detection of up to 100 analytes simultaneously.

Plasma samples were diluted in ratio 1:10 with Stabilguard buffer. All reagents and samples were adjusted to a room temperature before beginning of assay. Coupled beads should be vortexed 10 s and sonicated 10 s before adding to a master mix.

For the first incubation 50 μl diluted plasma sample was loaded on the plate together with 50 μl of bead mixture (containing 100 beads per μl for each region) and incubated at room temperature in the dark for 1 hour or overnight at +4 °C on the shaker. After incubation the plate was placed on the magnet for 60 s and aspirated followed by 3 washes with PBS.

For the second incubation beads were resuspended in 50 μl of Stabilguard and 50 μl of appropriate secondary antibody mixture was added. IgG was detected using biotinylated protein G (BPG) as a secondary antibody. For IgM antibody detection biotinylated anti-human IgM (affinity purified, μ-chain specific, Sigma-Aldrich cat. Nr. B1140) was used as a secondary antibody. Secondary antibodies were dissolved in StabilGuard to a concentration of 4 μg/ml. Plate was incubated in the dark with gentle rotation for 30 min. After the incubation the washing procedure was performed as described above. For the last incubation beads were resuspended in 50 μl of Stabilguard and 50 μl of streptavidin-phycoerythrin (SA-PE) (Molecular Probes, Leiden, The Netherlands) was added, diluted in Stabilguard buffer to 4 μg/ml. Beads were incubated for 15 min in the dark with gentle agitation. Before reading the plate, it was washed two times with PBS and beads were resuspended in 50 μl of Stabilguard.

### Non-denaturing elution (“neutralisation” assay) and crossreactivity test with human recombinant GPCRs

Beads with coupled immunogenic synthetic peptides were incubated with plasma samples pool overnight at 4°C in the dark. After incubation plate was placed on the magnet for 60 seconds and aspirated followed by 3 washes with PBS.

Beads were suspended in 100 μl of Stabilguard and divided into two samples by 50 μl each. First samples (50 μl) were used for antibody detection by SMIA as described previously, while their duplicates (second samples - 50 μl) were used for “neutralisation” assay by using Glycine buffer elution. This procedure was done for peptides showing immunogenicity for both IgG and IgM class antibodies.

20 μl of Elution Buffer (50 mM Glycine pH 2.8) was added to the appropriate bead samples and gently pipetted to resuspend. Samples were incubated with rotation for 2 min at room temperature to dissociate Ag-Ab the complex. After the incubation plate was placed on the magnet for 60 seconds and the supernatant containing dissociated antibodies was removed and placed in a new plate. The pH of the eluate can be adjusted by adding 1 M Tris, pH 7.5 and used for following incubation in crossreactivity test.

Beads after elution were resuspended in 50 μL of Stabilguard buffer and used for SMIA to test changes in MFI comparing results from the run with first samples (same samples but without elution buffer treatment).

Eluate acquired from neutralisation was incubated with xMag beads coupled with human recombinant GCPRs (CCR1, CCR3 and CCR5) and used in SMIA to test possible crossreactivity of peptide Ab acquired from elution.

## Results

### SMIA results of synthetic peptide specific IgG and IgM antibodies

All 30 synthetic peptides were tested for immunogenicity in 62 AIT patients’ plasma samples. Blank values and uncoupled beads were included in each SMIA run to exclude nonspecific antibody binding. Also in each coupling procedure, protein G or human IgM antibodies (depending on target class antibody) were coupled to one of the bead’s region ensure positive covalent binding of antigens during carbodiimide reaction and included in the followed SMIA run.

Of the 30 peptides tested on IgG immunogenicity, the strongest median fluorescent signal (MFI 264.8; IQR: 227.5 - 314.5) was shown by HHV-6_GPCR28 peptide, which was derived from the amino acid sequence of the HHV-6 U51 protein. Figure 2 shows the results obtained, with eight peptides with the highest MFI values coloured in red, which were used for further studies to assess cross-reactivity. Of the eight most immunogenic peptides, five were derived from the amino acid sequence of the U12 protein and three from the U51 protein.

**Figure 2.**
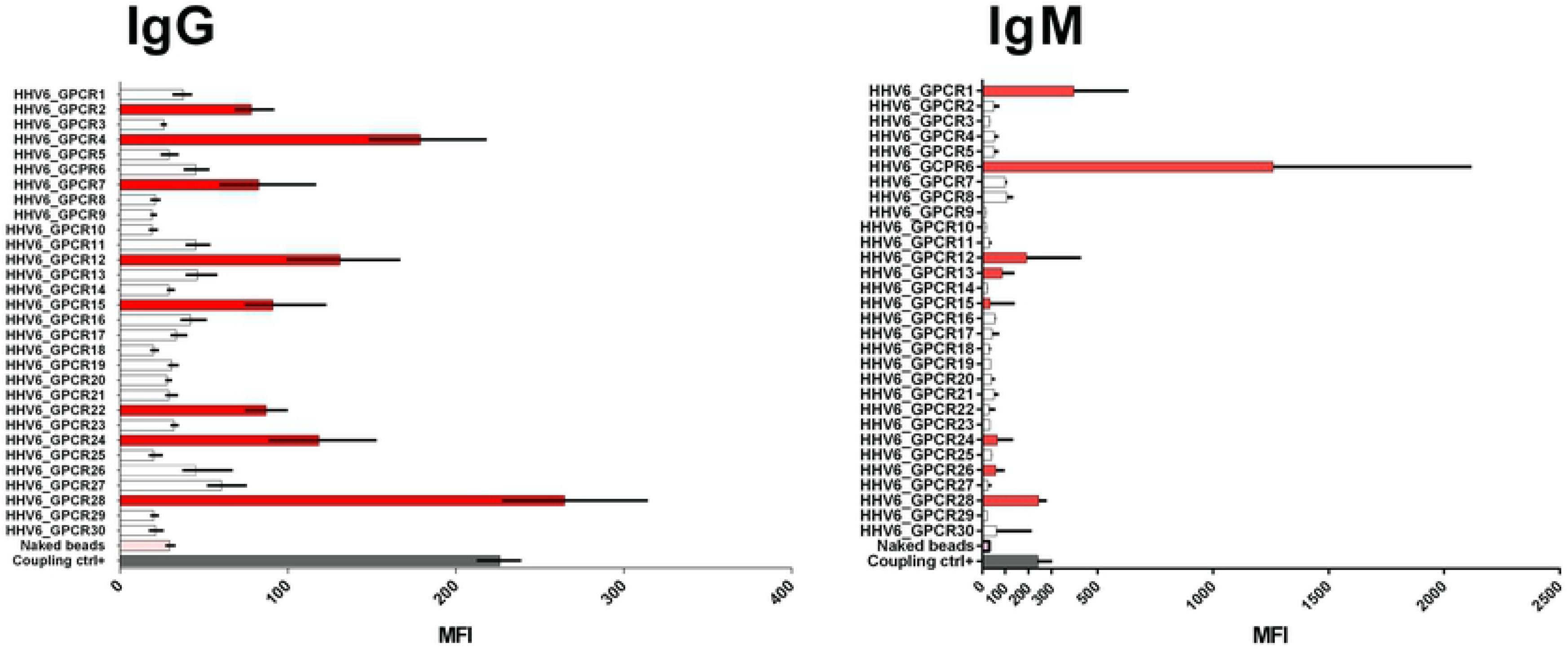
SMIA results on 30 synthetic peptides immunogenicity showing MFI values for IgG and IgM class antibodies

Of the 30 peptides tested on IgM immunogenicity, the strongest median fluorescent signal (MFI 1259; IQR: 440 - 2119) was shown by HHV-6_GPCR6 peptide derived from HHV-6 U12 protein. Figure 2 shows the results obtained, with eight peptides with the highest MFI values coloured in red, which were used for further studies to assess cross-reactivity. Of the eight most immunogenic peptides, five were derived from the amino acid sequence of the U12 protein and three from the U51 protein.

Screening of synthetic peptides immunogenicity showed higher MFI values for IgM.

### SMIA results on human recombinant protein CCR1, 3 and 5 specific IgG and IgM antibodies

Three human recombinant proteins CCR1, 3 and 5 were tested for IgG and IgM antibodies in patient plasma to find out possible presence of auto-antibodies against human GPCRs. The highest MFI values for IgG antibodies was observed in the case of CCR1 (MFI 47.3, IQR: 32 - 80), the obtained MFI values differed statistically significantly from the MFI values for unconjugated beads (p <0.0001) (Figure 3 IgG).

**Figure 3.**
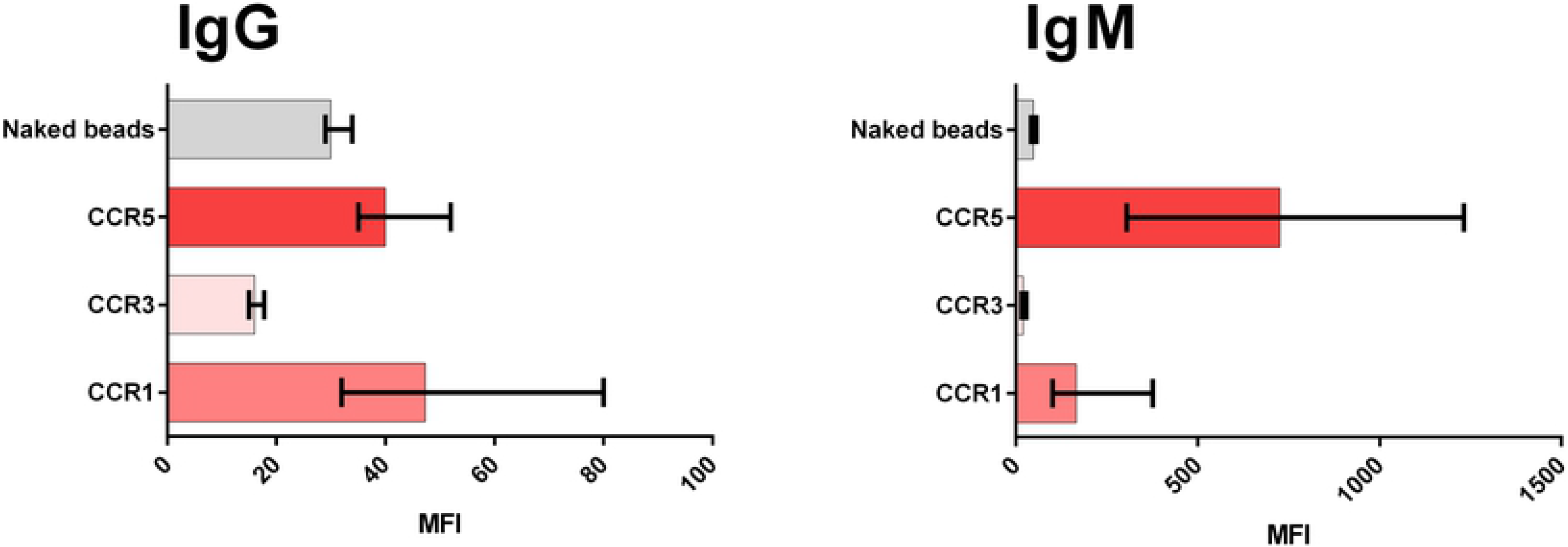
Human recombinant protein (CCR1, 3 and 5) specific antibodies (IgG and IgM)

The highest MFI values for IgM antibodies were for CCR5 (MFI 726, IQR: 304.6 - 1232), the obtained MFI values differed statistically significantly from the MFI values for unconjugated beads (p = 0.002) (Figure 3 IgM). In general, MFI values for recombinant proteins on IgM antibodies were significantly higher, with MFI values ranging to 80 in IgG assays but greater than 1200 MFI in IgM assays.

### Neutralisation and crossreactivity assay

Comparison of the same samples’ MFI values with and without elution (EL) showed not significant or no changes at all (Figure 4). This could indicate the absences of true IgG class antibodies or the presence of small amounts of antibodies, which could be none specific.

**Figure 4.**
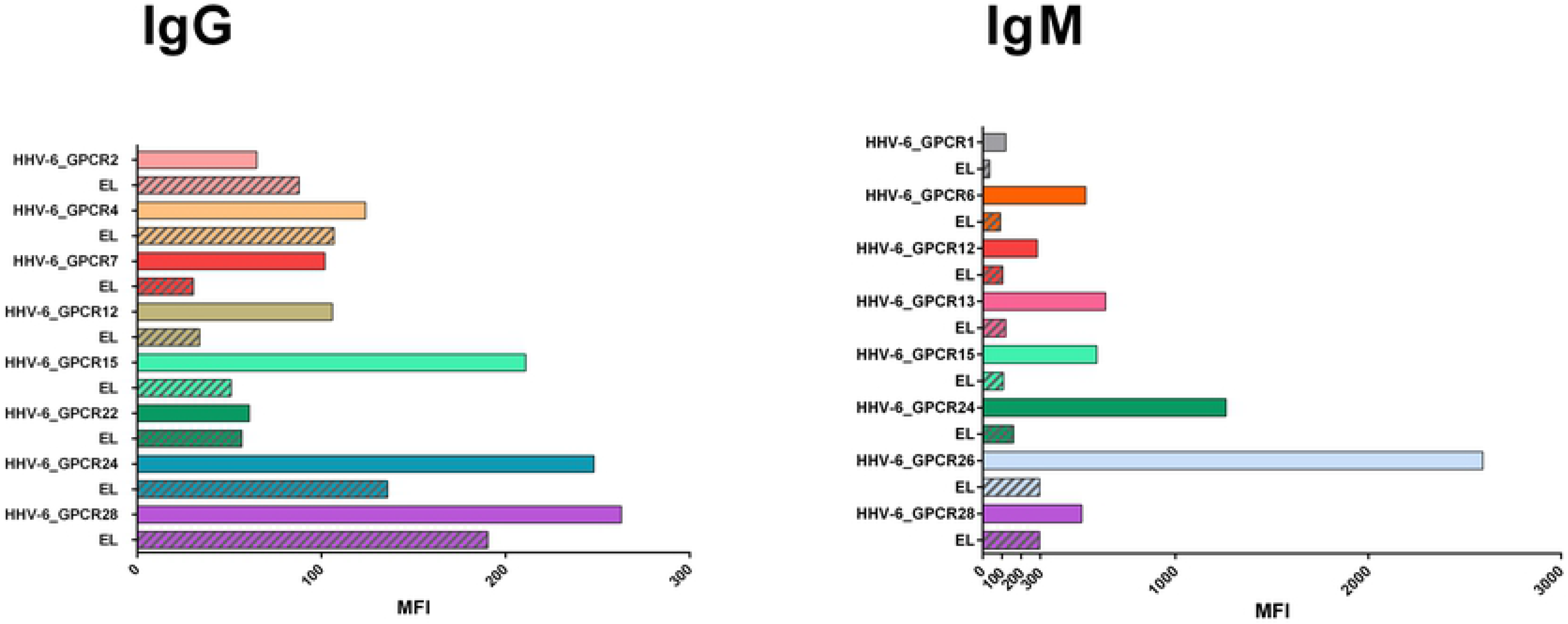
Comparison of plasma pool samples’ MFI values without and with elution (EL) of potential IgG and IgM class antibodies.

In contrast to IgG elution results, the same samples’ MFI values with and without elution (EL) of potential IgM class antibodies showed significant signal change (Figure 4). This could indicate that the immune response to the synthetic peptides is predominantly mediated by IgM class antibodies, which is also supported by the initially reported higher MFI values during the peptide immunogenicity assessment.

Eluated peptide specific antibodies were combined in four mixtures depending on antibody class and the viral protein from which the synthetic peptide was designed. Antibody mixtures were incubated in duplicate with beads coupled with human recombinant CCR1, 3 and 5, “none template controls” (NTC) were added. Obtained MFI values after incubation were compared with MFI values from NTC. No significant MFI value differences indicative of antibody crossreactivity were found for any of the antibody mixtures incubated with any of the human CCRs (Figure 5).

**Figure 5.**
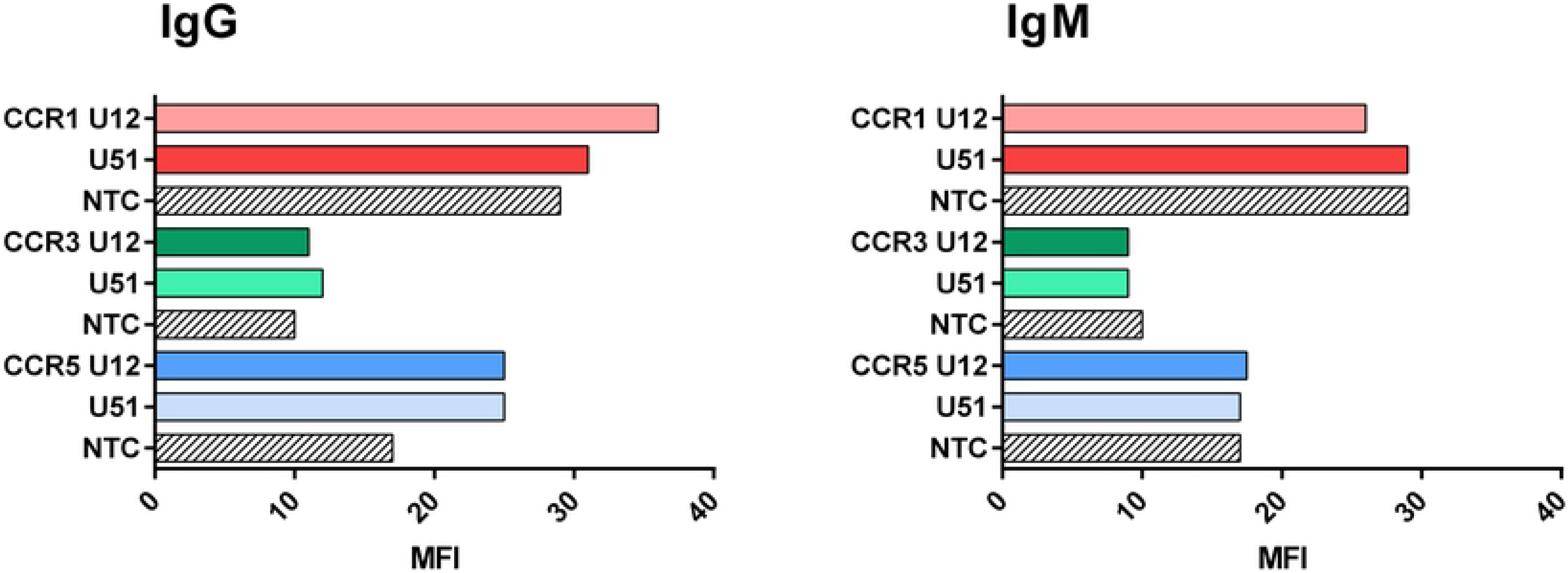
Crossreactivity assay results for IgG and IgM class antibodies.

## Discussion

HHV-6 is widely distributed in the general population. The primary infection usually occurs in the early years of life and remains latent throughout life [15]. HHV-6 has immunomodulating properties and has been shown to contribute to several autoimmune disorders, including autoimmune hemolytic anemia/neutropenia, autoimmune acute hepatitis, and multiple sclerosis [3–5]. Previous studies have showed high frequency of HHV-6 genomic sequence detected in thyroid gland samples obtained from patients with AIT [6,7]. In our latest manuscript, we show high frequency of HHV-6 U12 and/or U51 genes’ mRNA presence in thyroid gland tissues acquired from AIT patients [16]. Proteins encoded by these genes are structurally similar to cellular G protein-coupled receptors like human CCR1, CCR3 and CCR5 [10,12]. *In vitro* HHV-6 GPCRs may interact with the cytokine signalling pathway, leading to down-regulation of RANTES [14,17]. Also, our recent article shows significantly lower RANTES levels in AIT patients’ peripheral blood plasma in comparison with blood donors (median 150.3 [IQR: 71.6–418.2] pg/mL vs median 1359.0 [IQR: 844.2– 2596.0] pg/mL, respectively) [16]. Therefore, these viral proteins could be involved in autoimmunity development directly by interfering with RANTES signalling pathway and/or indirectly by mimicking cellular GPCRs. Due to the absence of on the market available HHV-6 U12/U51 proteins and difficulty in acquirement of recombinant HHV-6 U12/U51 proteins and antibodies to them, it was decided to use synthetic peptides designed from viral proteins’ amino acids. Use of synthetic peptides has its pros and cons. The main pro - they are cheap to produce and they can contain specific linear epitopes which could be used for specific diagnostic or for acquirement of specific antibodies. From the cons - the absence of full immunogenic information, for example, structural epitopes.

Synthetic peptides used in this study were designed based on analysis of two parameters - conservative regions from alignment with potential homologs (CCR1, CCR3 and CCR5) and predicted epitopes (using Bepipred Linear Epitope Prediction and Kolaskar & Tongaonkar algorithms). Such approach was chosen to increase chance of acquiring immunogenic peptides and meant to be more cost efficient for the implementation. However, this approach did not guarantee the possibility of identifying full linear epitopes. To fully describe linear epitopes further research should be done using overlapping amino sequences of peptides which were immunogenic in this study.

The most important finding of this study was the detected presence IgM antibodies against synthetic peptides designed both from HHV-6 U12 and U51 proteins’ amino acids, as well as, against human recombinant CCR1 and CCR5 proteins. This means that AIT patients have an constant immune response to hosts’ CCRs and to HHV-6 homologs (U12 and U51). This could indicate on possible involvement of HHV-6 in autoimmunity development as trigger factor.

Although, antibodies specific for synthetic peptides did not cross-react with human recombinant proteins, the possibility of cross-reactive autoantibodies specific for structural epitopes cannot be excluded. The structure of HHV-6 U12 and U51 proteins has not been elucidated. This makes it hard to find potentially immunogenic peptides due to the fact that their location in a protein in its native folding state is not known. Although there are softwares that help predict 3D structures of proteins they are only based on homology to other proteins. Further analysis of the peptides that showed highest MFI values in the immunogenicity assessment, showed that they most likely are located in the transmembrane part of the proteins, which could mean that it is physically impossible for the antibodies to bind even if they were cross-reactive.

## Acknowledgments

Supported by Project No.1.1.1.2/VIAA/1/16/202,Agreement No. 9.-14.5/257 “Human herpes virus-6 involvement in development of autoimmune thyroiditis”

